# Evaluation of UV-A LED technology on the reduction of spiked aflatoxin B_1_ and aflatoxin M_1_ in whole milk: toxicity analysis using liver hepatocellular cells

**DOI:** 10.1101/2021.03.14.435353

**Authors:** Anjali H. Kurup, Ankit Patras, Brahmaiah Pendyala, Matthew J. Vergne, Rishipal R. Bansode

**Affiliations:** Food Biosciences and Technology Program, Department of Agricultural and Environmental Sciences, Tennessee State University, Nashville, 37209, TN, USA; Department of Pharmaceutical Sciences and Department of Chemistry and Biochemistry, Lipscomb university, Nashville, 37204, TN, USA; Center for Excellence in Post-Harvest Technologies, North Carolina Research Campus, North Carolina A&T State University, Kannapolis, NC, USA

**Author notes:** Correspondence: Co-Corresponding authors: Ankit Patras, Ph.D., Associate Professor, Brahmaiah Pendyala, Ph.D. Research Scientist, /6018.

**Keywords:** aflatoxins, degradation kinetics, mechanism, photodegraded products, cytotoxicity

## Abstract

The effectiveness of a UV-A light emitting diode system (LED) to reduce the concentrations of aflatoxin B_1_, aflatoxin M_1_ (AFB_1_, AFM_1_) in whole milk (WM) was investigated. Irradiation experiments were conducted using an LED system operating at 365 nm. Known concentrations of aflatoxins were spiked in WM and irradiated at quantified UV doses which was calculated based on the average volumetric intensity. LC-MS/MS product ion scans were used to identify and semi-quantify photodegraded products of AFB_1_ and AFM_1_. It was observed that UV irradiation significantly reduced aflatoxins in WM, p<0.05. In comparison to control, the maximum UV-A exposure reduced AFB_1_ and AFM_1_ concentrations to 78.2 ± 2.36 % (at 836 mJ/cm^2^) and 65.7 ± 1.65% (at 857 mJ/cm^2^), respectively. In cell culture studies, our results demonstrated that the increase of UV-A dosage decreased the aflatoxins-induced cytotoxicity in HepG2 cells, and no significant aflatoxin-induced cytotoxicity was observed at highest given UV-A irradiation of 777 (AFB_1_), 838 (AFM_1_), and 746 (total AFs) mJ/cm^2^. Sensory quality of product, cytotoxicity, and mutagenicity of UV exposed aflatoxins in WM using animal models is warranted in the future.

## 1. Introduction

Fungal species that feeds on organic matter produces low molecular weight toxic secondary metabolites known as mycotoxins, that contaminate agriculture products and exert toxic effects on humans. It was estimated that 25 % of the world’s crops were contaminated by mold and fungal growth, in particular cereals, nuts, and oilseeds (Marin et al., 2013; Pandya & Arade, 2016). Foods of animal origin, such as milk and milk derivatives, eggs, and meat also possess a major threat of mycotoxicosis whose major source is from contaminated animal fodder (Bryden, 2012). The exposure of these chemical contaminants on humans can be through consumption of infected food and, in animals due to ingestion of infected animal feeds or by transfer of the toxins to their offspring during lactation or to other species through consumption of infected animal products like milk, meat, and eggs (Battacone et al., 2005).

Aflatoxins (AFs) derived from *Aspergillus flavus* and *Aspergillus parasiticus* have been among the most poisonous mycotoxins (WHO, 2018). The most frequent and cancerous members of the family of AFs as considered by IARC (The International Agency for Research on Cancer) were found to be aflatoxin B_1_ (AFB_1_) and its principal hydroxylated metabolite which is aflatoxin M_1_ (AFM_1_), both as a group I carcinogen (Bedard & Massey, 2006; Dai et al., 2017). Concerning the toxicity of AFM_1_, this toxin which was initially listed by IARC as a human carcinogenic agent in Group 2B (Organization & Cancer, 1993) has been reclassified as a Group 1 carcinogenic agent (Humans & Cancer, 2002). The fungal infections followed by subsequent production and accumulation of AFs occurs from the field itself during harvesting and continues even during storage. The carcinogenic risk assessment study of AFs by Cui et al., (2020) from post-fermented tea is evidence for this accumulation during storage. Direct or indirect exposure to these toxins gives off toxic, carcinogenic, mutagenic, teratogenic, and immunosuppressive effects on animals and humans (Arenas-Huertero et al., 2019). Studies have shown that life-threatening aflatoxicosis primes genotoxicity, meaning it can damage DNA and cause cancer in animal species (Attia & Harisa, 2016). The enzyme hepatic P450 cytochrome monooxygenase metabolizes AFB_1_ to reactive oxygen metabolites (AFM_1_, AFP_1_, AFQ_1_, and AFB_2a_) which once gets activated in the liver interacts with DNA and protein thus leads to growth retardation, immunosuppression, genotoxicity, and hepatotoxicity in the host (Kumar et al., 2017; Rushing and Selim., 2019). AFs were documented as the most potent congener, around 4.5 billion people in developing countries were infected with a daily toxin intake of 30 to 450 µg/L (Daniel et al., 2011). In the human food chain, milk is one of the most important sources of aflatoxins. While both M_1_ and B_1_ aflatoxins can be found in milk, the former is ten times more prevalent. From a public health standpoint, exposure to aflatoxin M_1_ through milk consumption has always been a greater concern than other aflatoxins due to the potential quantity of milk consumption as well as the vulnerable population (Arthur, 2012). World milk production has increased by more than 59 percent in the last three decades, from 530 million tonnes in 1988 to 843 million tonnes in 2018 (FAO, 2021). Hence, milk is considered as one of the high-risk foods that necessitates immediate action in turning out aflatoxin free.

Various federal agencies have set-up threshold levels (permissible limit) for mycotoxins in milk and milk products. For example, the United States Food and Drug Administration [USFDA] limits the total AFs levels to not more than 20 µg/kg whereas the European Commission (EC) limits to 4 µg/kg for food and feed meant for dairy consumption ((US Food and Drug Administration, 2000; Churchill et al., 2016). In the case of AFM_1_ alone, the regulatory limit for milk and milk products is 0.5 µg/kg (USFDA) and 0.05 µg/kg (EC) ((US Food and Drug Administration, 2000; Churchill et al., 2016). Moreover, there is a strong correlation between the concentration of AFB_1_ in animal feed and AFM_1_ contamination of mammal milk (Bakirci, 2001). AFB_1_ once ingested by animals gets bio-transformed to AFM_1_ within 24 hours and which may be present in all animal by-products derived from this intoxicated animal (Gürbay et al., 2006). The monohydroxy derivative AFM_1_ in the liver of lactating animals by the effect of cytochrome P 450 gets infused into milk in approximately 0.3 to 6.2 percent of the ingested AFB_1_ (Vaz et al., 2020). AFM_1_ concentration in 66 % of fresh dairy samples was reported higher than the European Union’s (0.05 µg/kg) maximum tolerance limit and 23 % higher than the US (0.5 µg/kg) maximum tolerance limit (Omar, 2016).

Traditional processing interventions are unable to degrade AFs due to their thermal stability and formation of acrid smoke at AFs decomposition temperature (> 237 °C) (Lewis et al., 2000). The greater binding property of AFM_1_ with casein increases the stability of toxin in curd. In cheese, the intensity of AFM_1_ is greatly related to the technology applied during cheese production but not on the ripening periods (Barbiroli et al., 2007). Infant formula milk, which is one of the most significant products made from milk is contaminated by AFM_1_ (Hooshfar et al., 2020). Though AFM_1_ is the only mycotoxin for which maximum residue limits has been established in milk, 0.4 µg/ L of AFB_1_ was also reported in heat treated milk (Udomkun et al., 2017). Hence, it is clear that AFs are milk contaminants that certainly creates a threat not only in milk but also in all derived dairy products (Oruc et al., 2006). All these findings reiterate the importance of degrading AFs in raw whole milk (WM) which can directly reduce the risk of aflatoxicosis in all milk products.

Latest advancements in science and engineering have demonstrated that light-based technologies hold considerable promise towards mycotoxins degradation (Udomkun et al., 2017). Various sources of ultraviolet light are commercially applied worldwide. Perhaps, these optical sources which are primarily lamps are unsustainable and have numerous drawbacks including footprint and production of mercury and other harmful waste (Hinds et al., 2020). As an alternative to UV mercury lamps, the application of light-emitting diodes (LEDs) has been under development (Shin et al., 2016). LEDs are an alternative optical source of UV light due to their lower energy requirements, longer life span, zero-waste production, and minimal heat generation (Bowker et al., 2011). LEDs are two-terminal semiconductor devices, which emit light under forward-bias conditions (Xu and Li., 2013).

Several investigative studies have been performed to test the effectiveness of LED in reducing the concentration of aflatoxins. Stanley et al., (2020) developed and tested a LED system against AFB_1_ and AFM_1_ in pure water. Although these tests were conducted in pure water, the authors demonstrated that an LED system at peak wave-length 365 nm can degrade aflatoxins effectively. Some studies focussed on oils and other food commodities also demonstrated an effective reduction of aflatoxins by UV energy. But the dose or fluence in most of these previous studies is incorrectly reported which makes it quite challenging to estimate the amount of UV delivered to the surface of the liquid sample of a specified volume. UV (220-400 nm) irradiation on peanut oil demonstrated complete AFB_1_ degradation in 30 mins but still did not report the dose delivered and optical property of the fluid (Liu et al., 2011). Similar gaps were observed in the study of Diao et al., (2015b) used 365nm UV light for AFB_1_ degradation in peanut oil.

The more sophisticated analysis includes the fluid optical properties and uses the volume-average intensity multiplied by residence time, which still ignores the impact of UV dose distribution (Chandra et al., 2017). For example, Mao et al., (2016) exposed peanut oil using UV-A irradiation and examined the aflatoxins by-products. The dose calculations did not incorporate the optical absorbance of the test solution. Without appropriate mixing, test fluid away from the optical source will receive a lower dose than that close to the free surface of the fluid (Chandra et al., 2017). The level of UV dose in a UV system will significantly depend on the distribution of dose values. Studies have also suggested UV-A as a better AF degradation tool compared to UVC due to its higher absorption coefficient (Diao et al., 2015a; Stanley et al., 2020).

The important questions related to the application of UV-A light in WM detoxifications includes: (a) optical properties characterization which includes separating absorption and reduced scattering coefficients including refractive index; (b) calculation of the irradiance and dose delivered by UV-A light a major problem in many UV studies, hence this study rectifies these problems by accounting for fluid optics and (c) understanding aflatoxins-induced cytotoxicity for liver cells in WM. Besides this study, there are no other reports in the scientific literature that evaluates cell cytotoxicity in UV treated WM. To the best of our knowledge, this is the first research study that has reported the ability of UV-A light to accelerate the degradation of aflatoxins in WM.

This research study investigates the effect of UV-A light on aflatoxins degradation in WM and determines the effectiveness of UV-A treatment of WM against aflatoxin-induced cytotoxicity for liver cells. Besides, our study also determined the degradation kinetics and the possible degradation mechanism of AFB_1_ and AFM_1_ in a highly scattering test fluid such as WM.

## 2. Materials and methods

### 2.1. Chemicals and reagents

Whole milk was obtained from a local Nashville, TN grocer and was immediately refrigerated at 4 °C. AFB_1_ and AFM_1_ were procured from LKT laboratories, Inc (St. Paul MN, USA). HPLC grade methanol, acetonitrile, and water were supplied by Sigma Aldrich (TN, USA). Supplies for the *in-vitro* cell cytotoxicity study such as Human hepatoma cells (HepG2; ATCC HB-8065), Eagel’s Minimum Essential Medium (EMEM; ATCC 30-2003), and fetal bovine serum (FBS; ATCC 30-2020) were purchased from American Type Culture Collection (ATCC, Manassas, VA). The XTT ([2,3-bis-(2methoxy-4-nitro-5-sulfophenyl)- 2H-tetrazolium-5carboxanilide) assay was purchased from Dojindo Molecular Technologies (Rockville, MD)

### 2.2. Optical measurements of fluids

For determining the optical attenuation properties of the test fluids, a double beam Cary 100 Spectrophotometer (Agilent Technology, Santa Clara, CA, USA) was used. The scattered light that is transmitted and reflected through the test sample in a thin quartz cuvette of defined path length was collected by the 6-inch single integration sphere (Lab sphere, DRA-CA-30, USA). An 8-degree reflectance port was used to measure the total reflectance. The device was initially calibrated to 100 percent and 0 percent baseline. Optical properties were measured as per the method described by (Ward et al., 2019). Ultraviolet transmittance (UV- A T %/cm), which is a calculation of the fraction of incident light transmitted along the 1 cm path-length of the sample, was measured using eq. 1. Transmittance measurements were described for scattering coefficient.

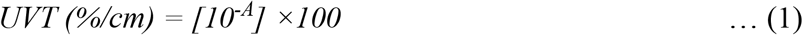

where A represents the absorbance (base_10_) of the test fluid at 365 nm for a 1 cm path-length.

### 2.3. UV-A exposure

A collimated beam system (Fig. 1) that houses a LED source (Irtronix, Torrence, CA, USA) producing irradiation at 365 nm was used for all the irradiations. To increase mixing, a 5 mL sample was stirred in a10 mL beakers (height: 3.2 cm; diameter: 1.83 cm) (Bolton & Linden, 2003; Chandra et al., 2017). All the optical parameters and UV dose calculations are described in our published paper (Stanley et al., 2020; Islam et al., 2016a; Islam et al., 2016b). Using a high sensitivity sensor (QE Pro series, Ocean Optics, Dunedin, FL, USA), the central irradiance of the UV-A LED system on the surface of the test solution was measured. The delivered UV dose in a stirred UV system was quantified (eq. 2) as the product of the volume average fluence rate in the sample and the exposure time by assuming adequate mixing of the sample by the stir bar.

**Figure 1:**
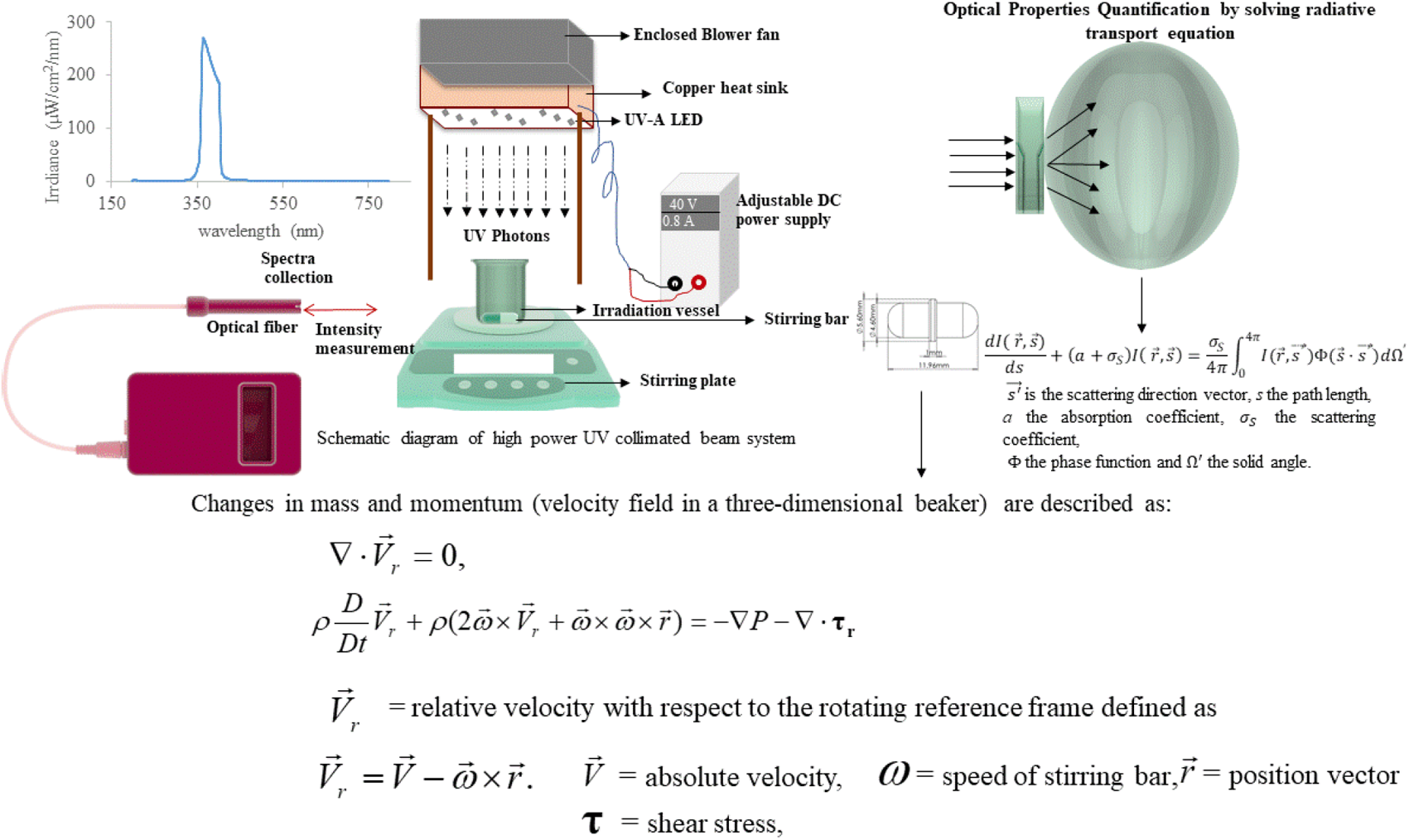
Experimental set up of UVA-LED system

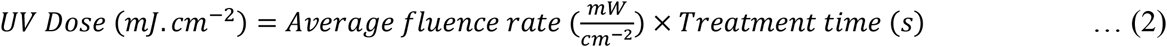

The average fluence rate was computed by the following eq. 3.

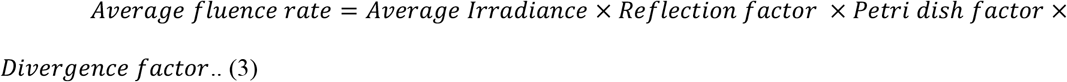

Average irradiance was calculated by computational Fluid Dynamics (CFD) simulation using ANSYS 12.1 software. The model considers the rotation of fluid (angular velocity) in the beaker exposed to incident UV incident radiation. The rotational speed of the magnetic stir bar was set to 240 rpm, which was the maximum speed that did not result in disturbances of the air-fluid interface. Modeling the UV system involved three sub-models: hydrodynamics; the intensity of the radiation field; and UV-reaction kinetics. The spatial UV irradiance was independent of the flow field, but a function of the optical properties (absorption and reduced scattering coefficient) of the WM, UV-output of the lamp, and distance to the LED source. After obtaining the velocity field by solving the transport equations, the Lagrangian Discrete Phase model was used to capture the trajectories of the fluid elements that follow the flow of the fluid. The other three correction factors i.e., reflection (RF) factor, Petri factor (PF), and divergence factor (DF) were accounted for in the average fluence rate calculations (Bolton et al., 2003).

AFB_1_ and AFM_1_ with initial concentrations of 1.5 µg/mL and 2.5 µg/mL respectively were spiked with the WM sample before irradiation. 5mL of WM with AFB_1_ was irradiated at 0, 35, 70, 139, 279, 557, and 836 mJ/cm^2^ providing uniform dose. Similarly, 5mL of WM with AFM_1_ was irradiated at 0, 36, 71, 143, 286, 571, and 857 mJ/cm^2^. UV doses and their corresponding exposure times are illustrated in table 1.

**Table 1:**
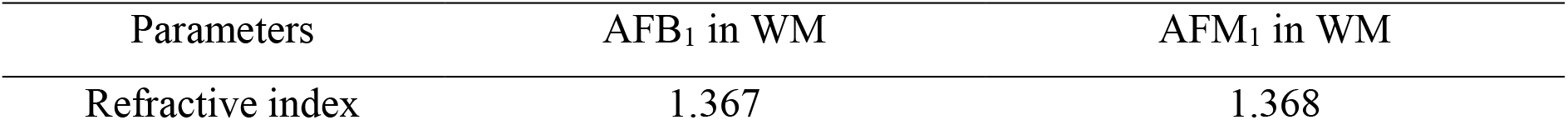
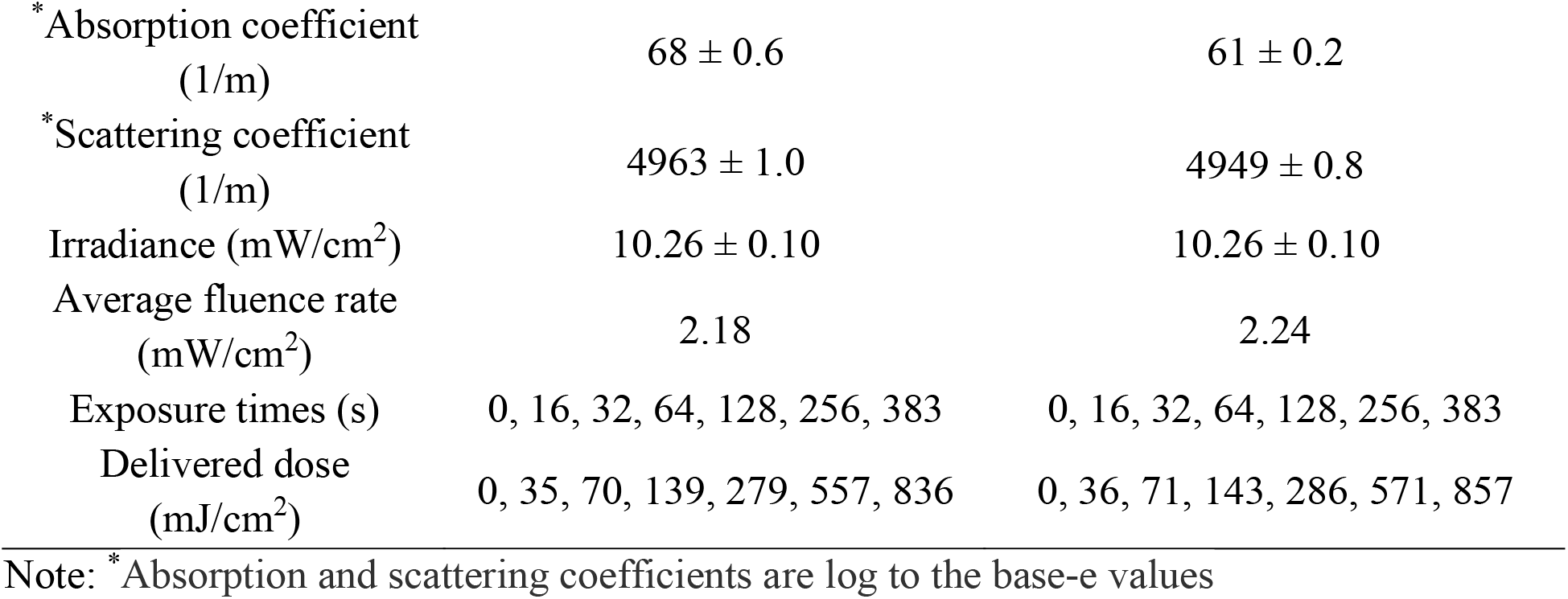
Optical properties and treatment parameters

### 2.4. Sample preparation for quantitative HPLC analysis

Irradiated and control samples were subjected to solid-phase extraction (SPE) to isolate the AF from WM. Strata-X 33 μm polymeric reversed-phase (Phenomenex®, USA) cartridges of 200 mg/3 mL were used for solid-phase extraction. 3 mL of the WM sample was loaded into pre-conditioned (with water followed by methanol) cartridges and was subjected to extraction under vacuum suction at a slow and steady phase. After the wash with 5 % methanol followed by 10 min vacuum drying, 3 mL of the methanolic extracts were eluted and the eluate was cleaned up with 0.2 μm PTFE filtration before HPLC analysis.

### 2.5. HPLC analysis of Aflatoxins

With slight modifications in the method described by Patras et al., (2017), the AFB_1_ & AFM_I_ content of the WM was measured. Briefly, 1 mL of the methanolic SPE extract of WM samples were injected into HPLC system (Shimadzu Scientific Instruments, Columbia, MD, USA). The sample carried by the mobile phase water/acetonitrile/methanol (120:50:30) with an isocratic flow of 1 mL/min, through a reversed-phase C_18_ column (Phenomenex, CA, USA) with configuration 150 mm × 4.6 mm 2.6 µm, maintained at 37 °C was eluted into Shimadzu RF20A fluorescence detector for an excitation and emission wavelengths set at 365 and 450 nm respectively. The standard calibration curve of AFB_1_ in methanol ranged between 0.25 to 1 µg/mL and that of AFM_1_ was from 0.5 to 2 µg/mL.

### 2.6. Evaluation of photodegraded AF products

A Shimadzu LCMS 8040 liquid chromatograph-mass spectrometry (LC-MS) system was used for the identification of AFB_1_, M_1_, and the respective degraded products were performed using selected ion monitoring. The injection volume was 5 μL, and the column used was the Phenomenex Kinetex C_18_ column (50 × 2.1 mm, 2.6 μm). As per (Patras et al., 2017) corresponding selected ion monitoring (SIM) MS methods for monitoring parent and degraded product-ions were identified. The mass-to-charge ratio of the degraded products of AFB_1_ (m/z 313) were m/z 303 and m/z 331. Similarly, for AFM_1_ (m/z 329), the mass-to-charge ratio of degraded products were m/z 347 and m/z 327. The dwell time for all SIM occurrences was 10 ms. The chromatographic peak regions of the treated samples and control samples had been calculated and compared with Shimadzu LabSolutions V5.89 software (Shimadzu, Columbia, MD, USA).

### 2.7. Kinetic modeling

GInaFiT tool was used to analyze the AF degradation kinetics results. GInaFiT, an add-in of Microsoft Excel was developed by (Geeraerd et al., 2005) as a depot of kinetic models, helps users determine the most precise model for fitting kinetics of degradation. A log-linear model developed by (Bigelow & Esty, 1920), were determined to be the best fit model to express the linear relationship between the dependent variable (concentration of AF in samples) and an independent variable (dose delivered). The model equation is given below,

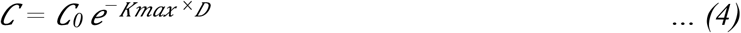

where *C* is the initial concentration of AF, *C*_*0*_ is the concentration of AF at dose *D, K*_*max*_ is the degradation rate constant and *D* is the UV dose delivered.

For identification purposes, the expression was reformulated as,

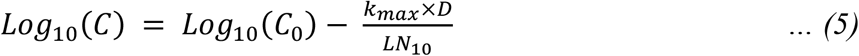

### 2.8. Effect of irradiated WM on human HepG2 cells

The study of cell cytotoxicity was conducted using a modified method as defined by Patras et al., (2017). HepG2 cells (ATCC; HB 8065) were seeded at 1.0 × 10^5^ cells per well in a 24-well plate containing Eagle’s minimum essential medium (EMEM) with 10 % (v/v) fetal bovine serum (FBS) and incubated for 24 hours at 37 ° C with 5% CO_2_. After 24 hours incubation, the cells were washed three times with PBS and serum starved in EMEM containing 1 % FBS overnight. Following overnight serum-starvation, the cells were exposed to 10 % (v/v) of untreated and treated test solutions at a final concentration of 3 μg/mL (each AFB_1_ and AFM_1_) and 6 μg/mL (total AFs). The cells were incubated for 24 hrs in the test solutions and cell toxicity measured using XTT (2,3-bis-(2methoxy-4-nitro-5-sulfophenyl)-2H-tetrazolium-5carboxanilide) assay as per the manufacturer’s protocol (Dojindo Molecular Technologies, Rockville, MD).

### 2.9. Data analysis

Three replicates for each treatment were exposed to the selected UV treatment. Each UV-A treatment was subjected to a balanced design in independent triplicates. Multiple comparison tests were carried out on one-way ANOVA with Tukey’s HSD to calculate UV-A effects in SAS computing statistical environment (SAS, 2016). Data are shown as a standard deviation from the mean. At 5 %, statistical significance was verified.

## 3. Results and discussion

### 3.1 Optical properties of whole milk

UV light (200 – 400 nm) transmission spectrum of whole milk was shown in Figure 2. The data show light transmission at UV-A region (315 – 400 nm) is higher than UV-C (200-280 nm) and UV-B (280 - 315 nm). Hence, UV-A has high penetration power in WM, thereby gradients in dose distribution throughout the sample are small, uniform dose distribution would be feasible with sufficient mixing. Since WM is a scattering fluid, both absorption and scattering coefficients were considered for the estimation of average light irradiance throughout the fluid. Absorption coefficients of WM (with AFB_1_ 68 ± 0.6 m^-1^ and with AFM_1_ 61 ± 0.2 m^-1^) were significantly lower than scattering coefficients (with AFB_1_ 4963 ± 1.0 m^-1^ and with AFM_1_ 4949 ± 0.8 m^-1^) (Table 1). The UV intensity of LED was measured as 10.26 mW/cm^2^, average volumetric fluence rate was calculated to be 2.18 (AFB_1_) and 2.24 (AFM_1_) mJ/cm^2^.

**Figure 2:**
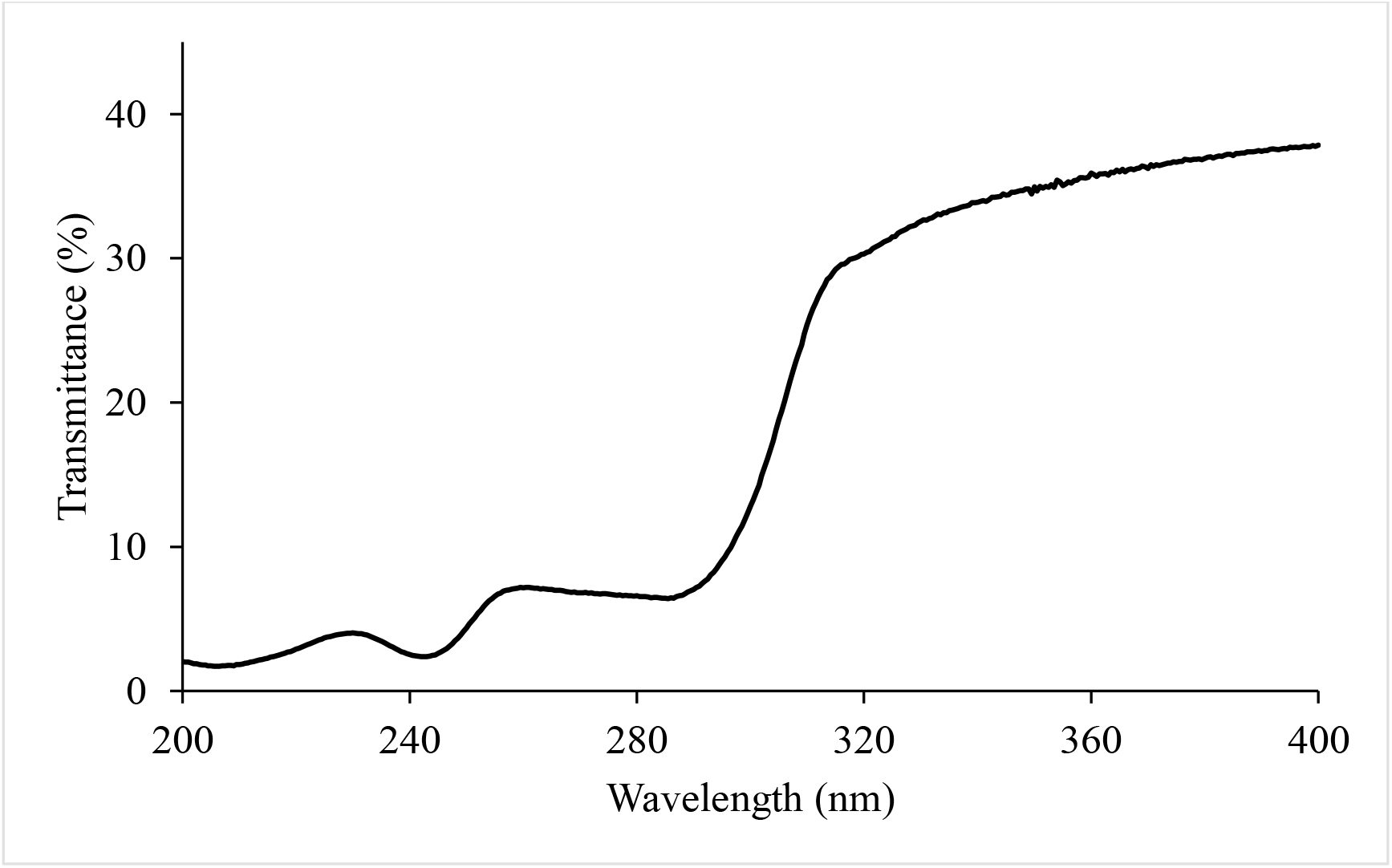
Transmission spectra of WM

### 3.2. Effect of UV-A irradiation on the degradation kinetics

Our previous reports demonstrated that AFB_1_ and AFM_1_ have strong absorption and acts as photosensitizers under UV-A irradiation (Stanley et al., 2020). AFB_1_ and AFM_1_ with initial concentration 1.5µg/mL and 2.5 µg/mL respectively in WM were subjected to UV treatment at doses ranging from 0 to 836 (AFB_1_) and 0 to 857 (AFM_1_) mJ/cm^2^. At doses 35, 70, 139, 279, 557, and 836 mJ/cm^2^ AFB_1_ reduction was observed to be 3.8 ± 1.33, 11.8 ± 0.65, 21.2 ± 1.66, 51.9 ± 1.13, 65.7 ± 4.51 and 78.2 ± 2.36 % respectively. In contrast, UV-A doses of 36, 71, 143, 286, 571, and 857 mJ/cm^2^ reduced AFM_1_ to 11.9 ± 1.85, 18.8 ± 1.04, 23.4 ± 2.11, 35.5 ± 2.30, 51.5 ± 3.93 and 65.7 ± 1.66% respectively. A log-linear model was fitted to the experimental data set. The goodness of fit and model parameters as illustrated in Table 2. Higher regression coefficients and low RMSE was observed.

**Table 2:**
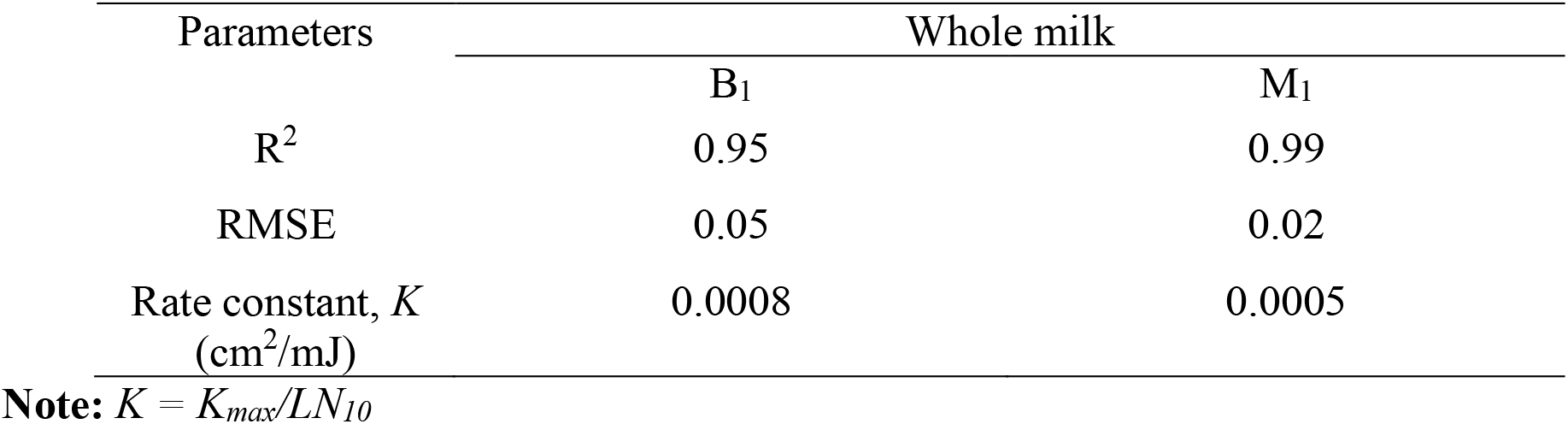
Model fitting and Goodness of fit

Fig. 3 illustrates the log-linear reduction of both AFs followed first-order kinetics with a maximum degradation of 78% (AFB_1_) and 65% (AFM_1_) at the highest dose of AFB_1_ (836 mJ/cm^2^) and AFM_1_ (857 mJ/cm^2^) respectively. The kinetic rate constant of AFB_1_ and AFM_1_ in WM was 0.0008 cm^2^/mJ and 0.0005 cm^2^/mJ respectively. The computed energy level was 5.44×10^−19^J and the average quantum yield (Table 3) of AFB_1_ was 1.59×10^−2^ and that of AFM_1_ was 1.14×10^−2^ in WM. In a different study, Stanley et al., (2020) reported AFM_1_ to be more susceptible to UV treatment in pure water with the degradation kinetic rate constant 1.6 times higher than that of AFB_1_. In contrast, our study demonstrated AFB_1_ to be highly sensitive as compared to AFM_1_ in WM. The degradation kinetic rate constant of B_1_ was 1.6 times that of M_1_. Quantum yield of AFM_1_ was 1.39 times less than AFB_1_, indicating AFM_1_ was relatively resistant to UV-A in comparison to AFB_1_. We hypothesize that this may be due to the presence of hydroxyl group in AFM_1_ which results in an increased affinity of M_1_ with milk proteins. Table 4 describes the predicted dose required to degrade AFM_1_ in WM below the FDA safety limit. For instance, if the initial AFM_1_ concentration in WM is 1.45 µg/L, to achieve the safety limit (0.5 µg/kg), a 65.7 % reduction is required which can be achieved with UV-A dose at 857 mJ/cm^2^ (assuming perfect mixing conditions).

**Table 3:**
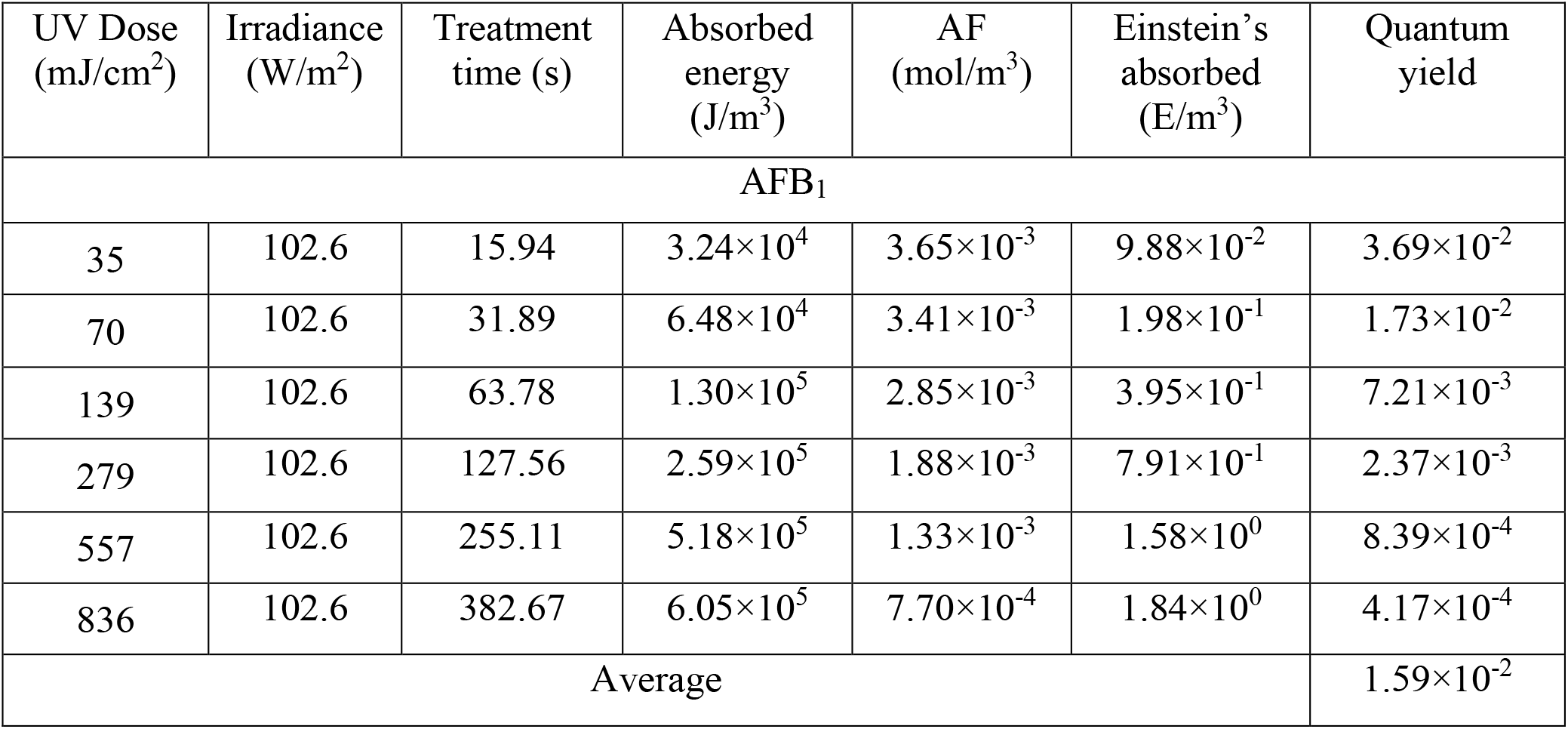
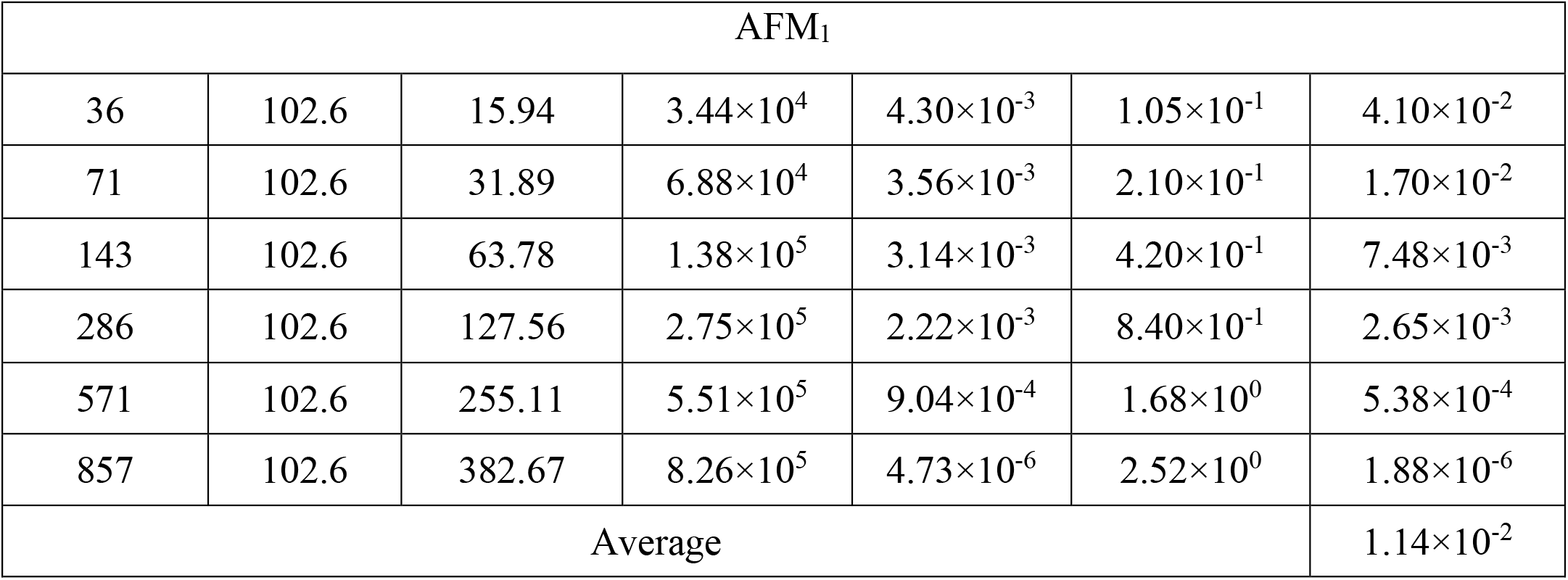
Quantum yield calculation of AFB_1_ and AFM_1_ in WM under UV-A radiation

**Table 4:** UV-A dose required to achieve safe limit conc. of AFM_1_ in WM.

| AFM<sub>1</sub> in WM<br>(µg/L) | Dose required<br>(mJ/cm<sup>2</sup>) to<br>achieve safety<br>limit | Reduction<br>achieved (%) | AFM<sub>1</sub> in<br>processed WM<br>(µg/L) |
| --- | --- | --- | --- |
| ≤ 0.56 | 36 | 11.9 | ≤ 0.5 |
| ≤ 0.61 | 71 | 18.8 | ≤ 0.5 |
| ≤ 0.65 | 143 | 23.4 | ≤ 0.5 |
| ≤ 0.77 | 286 | 35.5 | ≤ 0.5 |
| ≤ 1.03 | 571 | 51.5 | ≤ 0.5 |
| ≤ 1.45 | 857 | 65.7 | ≤ 0.5 |

Note: Based on FDA safety limit 0.5 µg/kg

**Figure 3:**
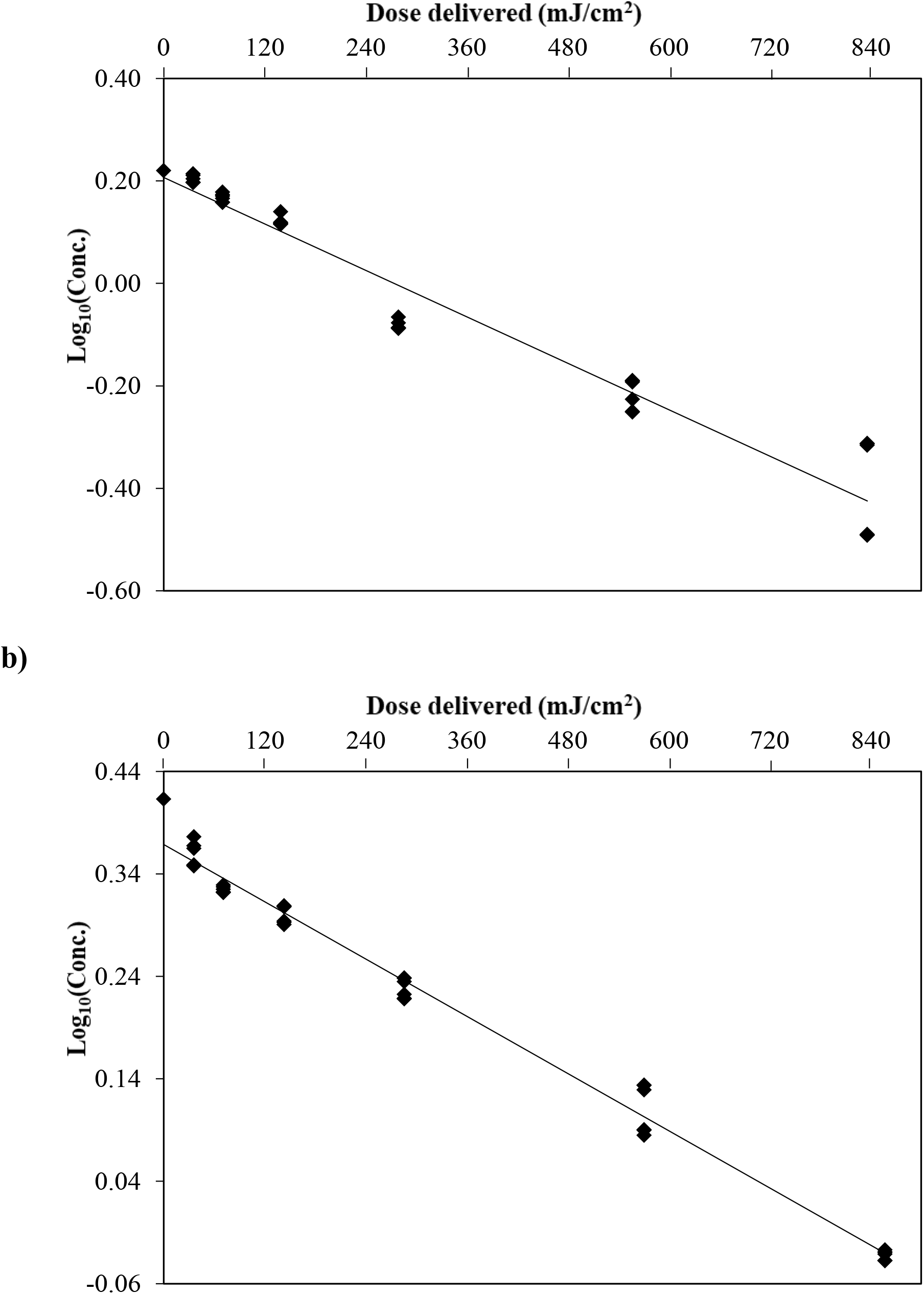
Degradation kinetics of (a) AFB_1_ and (b) AFM_1_ in WM

The photochemical fate of AFs with respect to per photon (UV) absorbed by targeted AFs (B_1_, B_2_, G_1_, G_2_, or M_1_) will be reflected as the change in compound concentration which can be quantified as quantum yield (Challis et al., 2014; Reina et al., 2018). It is evidenced that with increasing UV dose, the proportion of photodegradation also increases. Since UV irradiation produces the photons needed to pass electrons from the valence band to a semiconductor photocatalyst’s conduction band, and a photon’s energy is related to its wavelength, and the total energy input to a photocatalytic process depends on the intensity of light absorption. As more radiation falls on the catalyst surface, the rate of degradation increases forming more hydroxyl radicals (Pawar et al., 2018).

The formation of photosensitized singlet dioxygen can be ascribed as the basic degradation mechanism. UV light interacts with ground state oxygen and forms reactive oxygen species through a triplet excited state which is capable enough to detoxify AFB_1_ by detaching the lactone ring or furan ring, the primary sites of toxicity (Chandra et al., 2017; Stanley et al., 2020). Mao et al. (2016) also documented a similar UV detoxification mechanism on AFs. AFB_1_ functions as a photosensitizer under UV-A radiation and results in the formation of reactive oxygen species through either type I or type II mechanisms through the intervention of the triplet excited state. In type I mechanism electron transfer occurs from triplet AF to molecular oxygen, resulting in the formation of radical anion superoxide. Energy is transferred from triplet AF to molecular oxygen in the Type II mechanism, which contributes to singlet oxygen formation. These reactive species respond to AFs and cause the compounds to incur photolytic damage (Netto-Ferreira et al., 2011).

### 3.1. LCMS/MS for the analysis of photodegraded products of AFB_1_ and AFM_1_

Molecular formulas and elemental composition of degraded AFB_1_ and AFM_1_ products in WM samples were analyzed and predicted by LC-MS with a single ion monitoring system (SIM). WM spiked with a known concentration of AF without UV photolysis were used as a control sample. LC-MS peak areas for AFB_1_ and AFM_1_ decreased as compared to controls, and the peak areas of the degraded products increased significantly with an absolute increase in UV-A dose. As reference standard are not readily available for photodegraded product peaks, a quantitative approach is not feasible. For AFB_1_ and AFM_1_, representative SIM chromatograms are provided in Fig. 4. For the protonated adduct [M+ H]^+^, AFB_1_ and M_1_ showed strong ionization efficiency in positive ion mode with molecular base ion at m/z 313 (retention time, RT, 2.62 min) and m/z 329 (RT, 5.5 min) respectively. The protonated molecule was selected for monitoring in SIM because it is more intense than sodium adducts (Iram et al., 2015). As shown in Fig. 5, the structure of the degraded AFB_1_ and AFM_1_ products has been established and the potential pathway to degradation is proposed. The high-resolution mass chromatogram showed C_17_H_14_O_7_ (P1, m/z 331, RT 2.17 min) and C_16_H_14_O_6_ (P2, m/z 303, RT 2.26 min) as two product-ions of AFB_1_. Similarly, AFM_1_ also showed two degraded products, C_17_H_14_O_8_ (Q1, m/z 347, RT 4.18 min) and C_17_H_10_O_7_ (Q2, m/z 327, RT 5.37 min). The major fragmentation pathways of AFB_1_ with a double bond equivalence (DBE 12) were hydration and demethoxylation. The photolysis product P1 (DBE 11) was the result of hydration on the double bond of the terminal furan ring of AFB_1_ and has a comparatively lower toxic potential than AFB_1_. Product P2 (DBE 10) was the result of benzene side-chain hydration on the furan ring and demethylation. Hydration was reported as the major degradation pathway of AFM_1_ (DBE 12) into Q1 (DBE 11) and, Q2 (DBE 13) by adding the oxygen atom to the left furan ring double bond (Iram et al., 2015). The double bond in a furan terminal ring is known to be the most toxicological site of AFB_1_ with a lactone ring in the coumarin (Stanley et al., 2020). The biotransformation of AFB_1_ due to protein binding or formation of respective DNA adduct activates the leading active site for its toxic and carcinogenic function (Li & Liu, 2019). The double bond epoxidation results in the production of reactive oxygen species metabolite (AFB_1_-8,9-epoxides) (Benkerroum, 2020). It is seen from the literature that the toxicity of the photolysis products is decreased in contrast with the toxicity of AFB_1_ and AFM_1_ based on quantitative structure-activity relationships (Mao et al., 2016).

**Figure 4:**
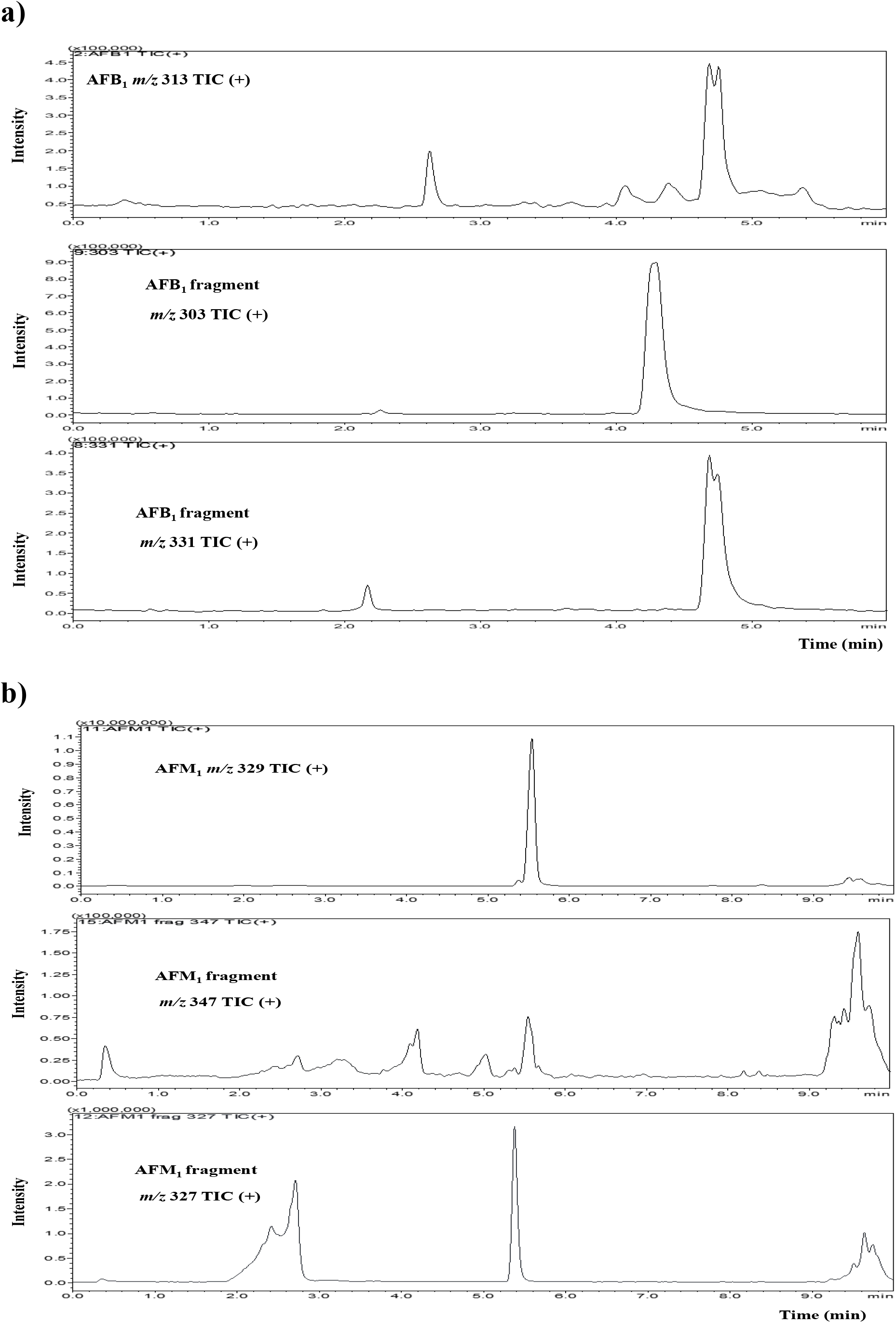
LCMS single ion monitoring (SIM) total ion chromatograms (TIC) of (a) AFB_1_ at 836 mJ/cm^2^ and (b) AFM_1_ (at 857 mJ/cm^2^)

**Figure 5:**
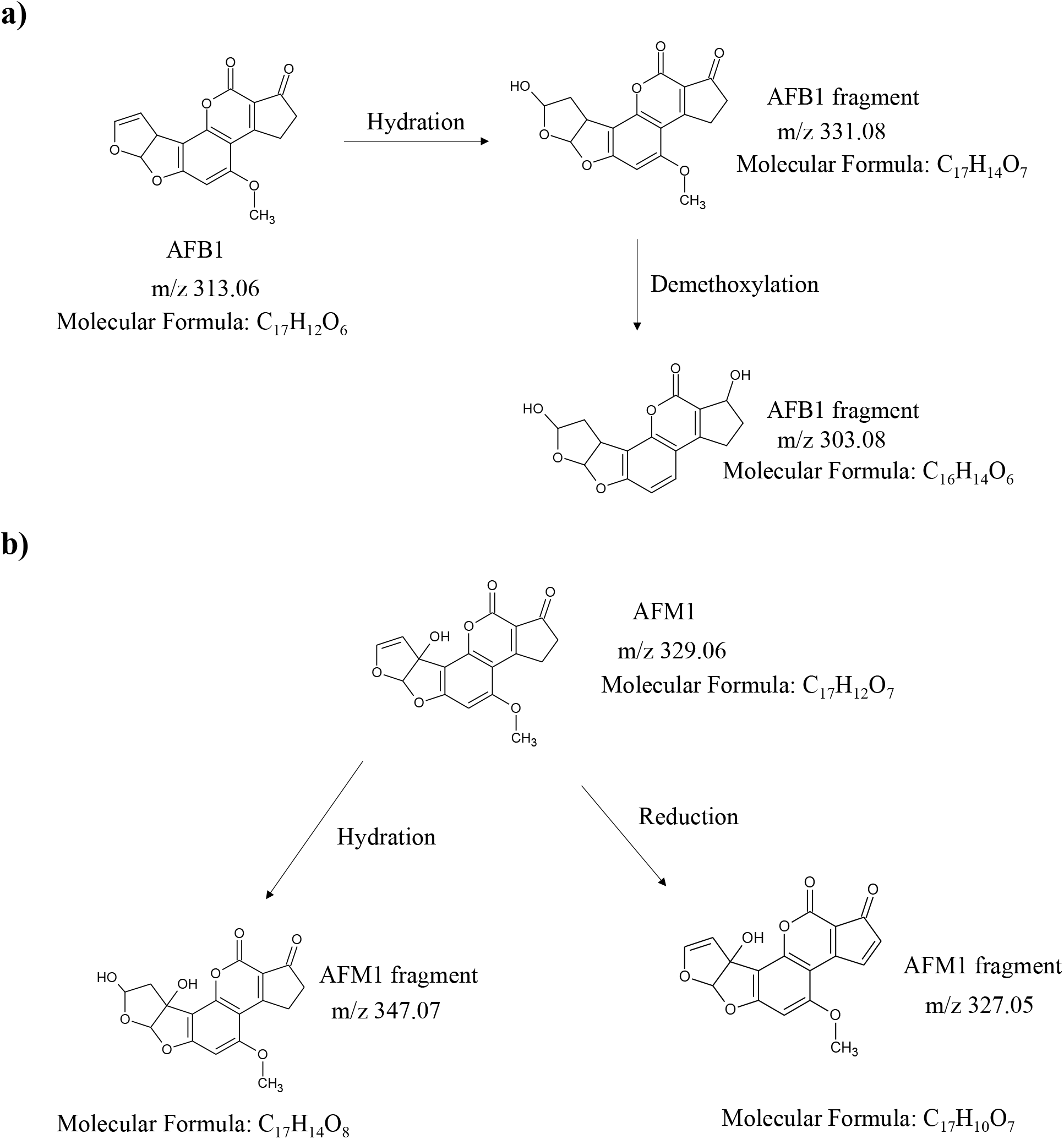
Proposed UV-A light degradation mechanism of (a) AFB_1_ and (b) AFM_1_ in WM

### 3.2. Effect of irradiated WM on human HepG2 cells

The cytotoxic mechanism of AFB_1_ is greatly associated with the generation of reactive oxygen species that damages DNA and cell membrane (Zhang et al., 2015; Zarev et al., 2020). The metabolism of AFs in many species under different conditions was widely studied through *in vivo* and *in vitro* analysis mainly on AFB_1_ which are easily absorbed and diffused into the mesenteric venous blood chiefly from the duodenum of the small intestine. Due to the reason that the liver has a functional link with the intestinal nutrients in the blood of the gut, the liver is the first organ affected by AF (Yunus et al., 2010). The cytochrome P450 coverts AFB_1_ to highly unstable reactive metabolite, AFB_1_-8,9 epoxide, which is the carcinogenic form, and then further converted to AFB_1_-dihydrodiol (Magan et al., 2010). Interaction of dihydrodiol and protein amino groups produces Schiff base adduct resulting in acute toxicity, while the chronic toxicity is caused by the combination of AFB_1_ with guanine moiety of DNA producing aflatoxin-N7-guanine adduct (Baertschi et al., 1989).

Since several studies have reported the toxic action of aflatoxin on the liver (Liu et al., 2014; Van Vleet et al., 2006), the photodegraded product in our study was investigated for cytotoxicity analysis against human HepG2 liver cells using XTT serum biochemical assays. Viability of the liver cells exposed to UV-A for 128 and 383 s with respective test fluids of AFB_1_ (calculated doses 259 and 777 mJ/cm^2^), AFM_1_ (calculated doses 279 and 838 mJ/cm^2^) and total AFs (calculated doses 249 and 746 mJ/cm^2^) in WM are presented in Fig. 6. This cytotoxic cell injury caused due to the free radical production of AF can be due to up-regulation of the expressions of caspase-3, caspase-9, CYP3A13, Bax, and p53 and down-regulation of the expression of TNF-α and Bcl-2 and their target proteins and thereby causing the oxidative stress damage in the cells (Peng et al., 2016).

**Figure 6:**
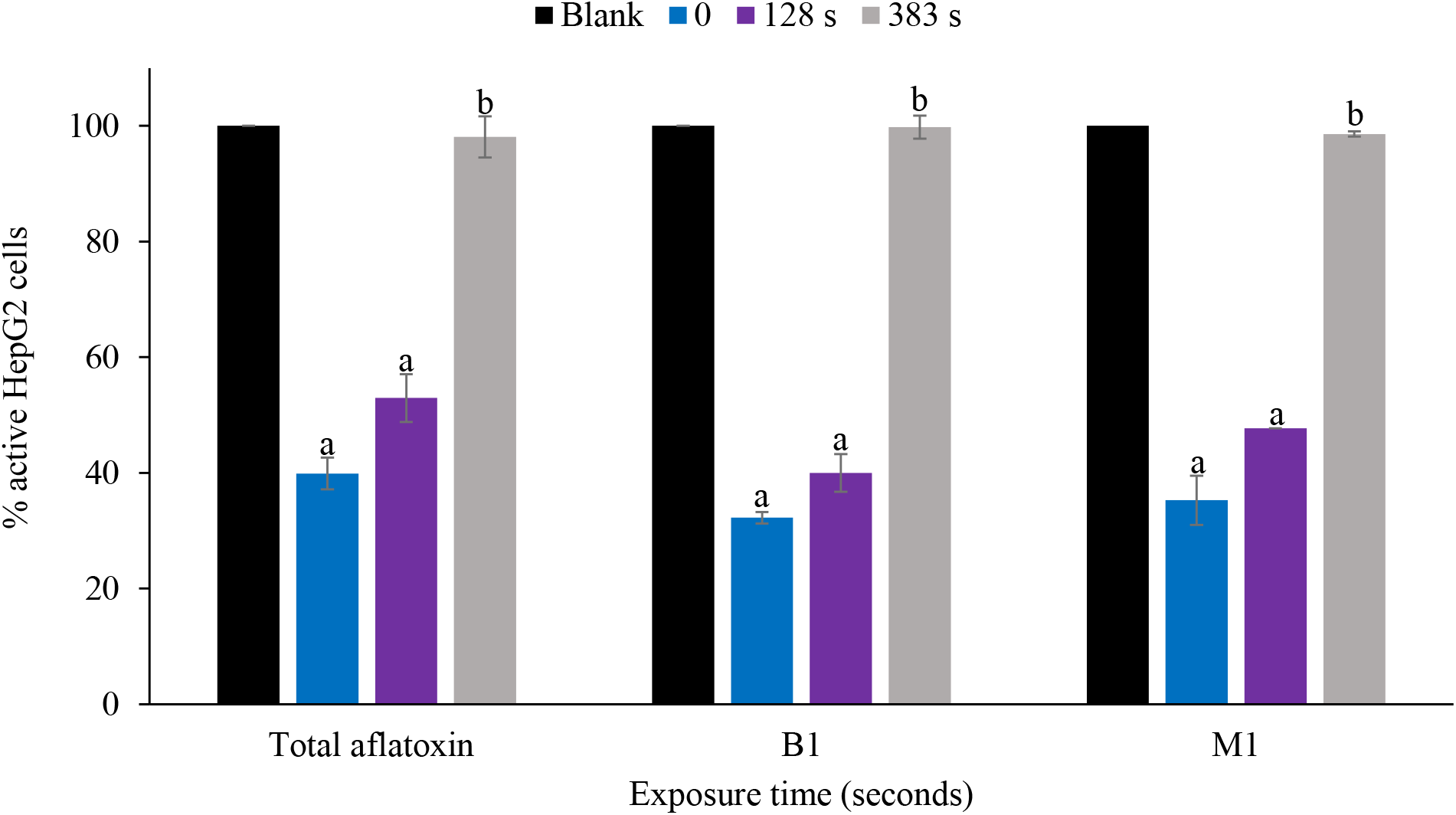
Cell viability evaluation after 24 h incubation **Note**: positive control (no AF, no UV irradiation); 0 (spiked with AF, no UV irradiation); dose delivered (mJ/cm^2^) at 128s-249 (total AFs), 259 (AFB1), 279 (AFM1); dose delivered (mJ/cm^2^) at 383s – 746 (total AFs), 777 (AFB1), 838 (AFM1). Data are presented as mean percentage ± SD of three separate experiments. Levels connected by different letters are significantly different at p < 0.05.

Three sets of experiments were conducted: HepG2 cells treated with 6 µg/mL of total AF (M_1_+B_1_), HepG2 cells treated with 3 µg/mL of B_1_, and HepG2 cells treated with 3 µg/mL of M_1_. A positive (no toxin) control was included for each experiment. Initially, 24-well plate with 100,000 HepG2 cells per well was incubated with the samples for 24 h and the relative % of active cells in the wells were measured. The relative percentage of viable cells as measured by XTT assay showed a linear trend with increasing UV dose delivered (Fig.6). In the untreated samples containing toxins, only 39.9 ± 2.76, 32.3 ± 0.99, and 35.3 ± 4.24 % of the hepatoma cells remained viable due to the impact of total AF, AFB_1_, and AFM_1_ toxicity respectively. However, UV treatment of 249 (total AFs), 259 (AFB_1_) and 279 (AFM_1_) mJ/cm^2^ significantly increased the cell viability to 52.9 ± 4.11, 40 ± 3.28, and 47.7 ± 0.01 % respectively. The highest dose delivered was quantified to be 746 (total AF), 777 (AFB_1_), and 838 (AFM_1_) mJ/cm^2^ which successfully achieved cell viability of 98.1 ± 3.55, 99.8 ± 2.01, and 98.6 ± 0.42 % respectively, showing a positive correlation between UV treatment and cell survival. There was no significant difference (p < 0.05) in the % relative cell viability between the control sample with no aflatoxins and UV treatment of aflatoxin samples that received a higher dose signifying the detoxification of WM under our experimental conditions. Therefore, the lack of significant toxicity in samples treated at these doses could be due to the presence of a very low concentration of AFB_1_ and AFM_1_ to induce toxicity in HepG2 cells within 24 hrs of exposure time or that the degradation products exhibited cell proliferation properties.

The cytotoxicity data signifies that the photodegraded products do not leave behind toxic metabolites that can decrease the cell viability, also it is clear no new hazardous contaminants are produced due to UV treatment in WM. Zhao et al., (2020) have shown that the AFB_1_ degradation products lacking lactone ring are either less toxic or exhibit growth-promoting effects. However, we are yet to evaluate the characteristics of the degradation compounds from the UV-treated samples from our experimental conditions. Our results are in accordance with some of other UV-related studies with no toxic effect on the irradiated i.e., coconut water (Bhullar et al., 2018), skim milk (Ward et al., 2019), and apple juice (Chandra et al., 2017). The HepG2 cytotoxicity study on UV irradiated ultrapure water also demonstrated no significant loss of the active cells (Stanley et al., 2020).

It was observed that the difference in cell viability between the samples treated and the controls were positively significant (p < 0.05). There was a substantial increase in the concentration of active HepG2 cells (p-value 0.002) with increasing dose from 0 to 777 (AFB_1_), 0 to 838 (AFM_1_), and 0 to 746 (total AFs) mJ/cm^2^, which demonstrates the efficiency of UV-A on aflatoxin degradation in WM.

## 4. Conclusion

Results from this study demonstrated that UV-A irradiation successfully reduced aflatoxins concentration in WM, suggesting that UV-A treatment is a plausible method to detoxify AFB_1_ and AFM_1_. This study confirms the importance of the quantifying optical properties, namely absorption and scattering values of highly opaque fluids to efficiently reduce aflatoxins. It was observed that more than 65% and 78% reduction of AFM_1_ and AFB_1_ respectively in WM can be achieved at 836 and 857 mJ/cm^2^ respectively, in challenging opaque fluids. Cell culture studies suggested that increasing UV-A dose in aflatoxin-spiked WM decreased the aflatoxin-induced cytotoxicity in Hep2 cells. Cell viability percentage increased significantly to 98.1% at maximum exposure dose. Further studies on understanding the degradation mechanism of aflatoxins by UV-A irradiation, sensory attributes of the product, assessing the cytotoxicity and mutagenicity of UV exposed WM using animal models, and evaluating the efficiency of the process on pilot scale is warranted.

## Acknowledgements

This project is funded under the Agriculture and Food Research Initiative (Food Safety Challenge Area), USDA, Award number; 2018-38821-27732.

**Note:** There are no conflicts to declare

